# Discovery of pseudobaptigenin synthase, completing the (-)-maackiain biosynthetic pathway

**DOI:** 10.1101/2025.11.18.689130

**Authors:** Lee Marie Raytek, Lan Liu, Stéphane Bayen, Mehran Dastmalchi

## Abstract

Pterocarpans are structurally complex defence compounds produced by legumes (Fabaceae). They are commonly associated with antimicrobial activity and thought to be synthesized *de novo* or accumulated in response to microbial pathogens. (-)-Maackiain is a lineage-specific pterocarpan detected in some legumes, including red clover (*Trifolium pratense*). The biosynthesis of (-)-maackiain involves a distinctive methylenedioxy bridge formation step, predicted to be catalyzed by a cytochrome P450. Specifically, this elusive P450 catalyzes the conversion of calycosin to pseudobaptigenin. We integrated metabolomic and transcriptomic datasets of red clover roots treated with the fungi, *Fusarium oxysporum* and *Phoma medicaginis*, to identify candidate P450 genes. Over 40 molecular features were characterized as (iso)flavonoid structures, including the highly abundant *O*-methylated isoflavones (formononetin and biochanin A), as well as their derivatives. Long infection with *P. medicaginis* resulted in significant increases in (-)-maackiain, trifolirhizin and other pterocarpans. Concurrently, fungal infections led to upregulation of core and specialized metabolism-related transcripts, including those encoding phenylpropanoid and (iso)flavonoid biosynthetic enzymes. Using weighted gene co-expression network analysis (WGCNA), variance-stabilized expression patterns, and enzyme-class phylogeny, we were able to curate five candidate cytochrome P450s for pseudobaptigenin synthase (PbS) activity, assayed in engineered yeast (*Saccharomyces cerevisiae*). One candidate P450 was capable of methylenedioxy bridge formation, converting calycosin to pseudobaptigenin and pratensein to 5-hydroxypseudobaptigenin. Therefore, it was renamed *T. pratense* pseudobaptigenin synthase (TpPbS/CYP76F319). The discovery of TpPbS facilitates the reconstruction of the complete (-)-maackiain biosynthetic pathway and the production of this pterocarpan chemistry at scale for health and agricultural applications.

**Significance statement:**

- The discovery of pseudobaptigenin synthase in red clover (CYP76F319), a P450 that catalyzes the formation of a methylenedioxy bridge to convert calycosin to pseudobaptigenin and pratensein to 5-hydroxypseudobaptigenin.
- The identification of enriched (iso)flavonoids and associated transcriptomic changes in red clover roots in response to two fungi with distinct infection lifestyles (hemibiotrophy and necrotrophy).

## INTRODUCTION

Plants are besieged by microbes in their environment, including pathogens, that seek to colonize their tissues and gain access to their resources. The fact that plants remain largely immune to these advances is remarkable. Beyond physical barriers, an aspect of this successful defence, or resistance, is raised by antimicrobial compounds that are synthesized and accumulate upon pathogen challenge, namely “phytoalexins.” The term, coined by Müller and Börger, and later refined by Paxton, stresses the *de novo* biosynthesis of these chemicals (1, 2). Legumes (Fabaceae) produce a large repertoire of lineage-specific phytoalexins, including a suite of compounds classed as pterocarpans. They have a characteristic benzofuran-benzopyran scaffold and their accumulation, particularly in the roots, has been associated with resistance against pathogens (3, 4).

The pterocarpan scaffold is derived from the legume-characteristic isoflavones, which are more constitutively accumulated. Isoflavonoids are a sub-class of flavonoids, where the shikimate-derived phenyl ring has migrated from the 2-to the 3-position of the chromone, a rare occurrence in nature (5). Structurally, pterocarpans are enantiomeric compounds, distinguished from other isoflavonoids by a pentacyclic ring closure at the O-11 position. Moreover, they are synthesized in a species-specific manner, exemplified by (-)-glycinol and the (-)-glyceollins (*Glycine*), (-)-phaseollins (*Phaseolus*), (-)-medicarpin (*Cicer*, *Glycyrrhiza*, and *Medicago*), and (+)-pisatin (*Pisum*) (6–10). The pterocarpans have been heavily implicated in defence against destructive crop pathogens, including the soil-borne oomycete *Phytophthora sojae* and the grey mold *Botrytis cinerea*, highlighting their potential utility as natural antimicrobial agrochemicals (3, 11).

Pterocarpan biosynthesis typically proceeds through a series of enzymatic steps downstream of the isoflavone aglycones, daidzein and genistein. For example, the formation of (-)-medicarpin starts with the *O-*methylated isoflavone formononetin (7-hydroxy-4′-methoxyisoflavone), which is 2′-hydroxylated by isoflavone 2′-hydroxylase (I2′H), followed by consecutive reductions catalyzed by isoflavone reductase (IFR) and 2′-hydroxyisoflavanone 4-reductase (I4R), respectively. This pathway is completed by pterocarpan synthase (PTS), which catalyzes a ring closure to generate (-)-medicarpin and the pterocarpan scaffold (10, 12–16).There are many parallel and overlapping branches to this canonical pathway, comprising a dense and diverse chemical landscape.

In red clover (*Trifolium pratense*), the highly abundant *O*-methylated species, formononetin and its 5-hydroxy counterpart, biochanin A (5,7-dihydroxy-4′-methoxyisoflavone), can undergo additional modifications before entering the core pterocarpan biosynthetic module described above. This branch begins with 3′-hydroxylation of formononetin by isoflavone 3′-hydroxylase (I3′H) yielding calycosin (17), which harbours adjacent hydroxyl and methoxy groups that can be enzymatically linked via a methylenedioxy bridge (MDB), producing pseudobaptigenin (Figure 1). The latter reaction has been described but the enzyme has not been identified (18). This new intermediate follows the same suite of reactions described above in (-)-medicarpin biosynthesis, i.e., 2′-hydroxylation, two successive reductions, and a ring closure to generate the pterocarpan (-)-maackiain. Likewise in a parallel route, biochanin A is 3′-hydroxylated to form pratensein, which can undergo MDB formation to 5-hydroxypseudobaptigenin (17, 18).

**Fig. 1.**
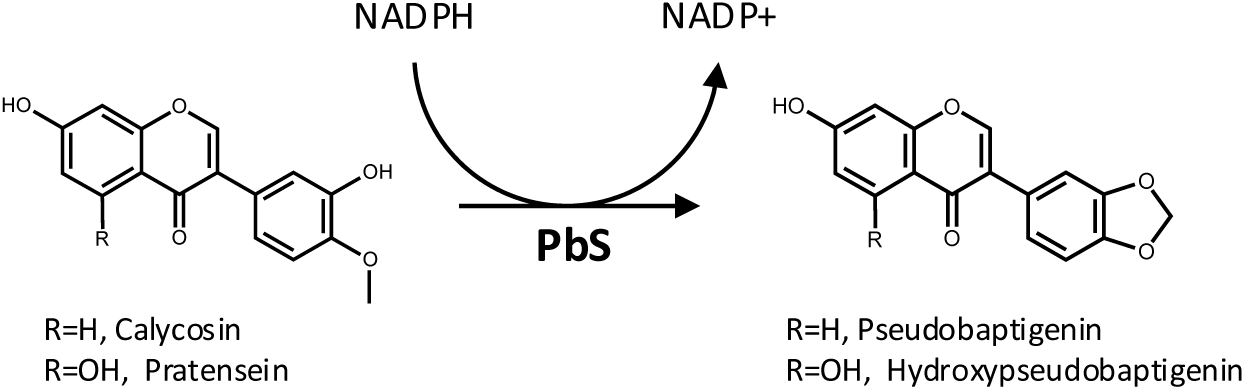
Methylenedioxy bridge formation by pseudobaptigenin synthase (PbS). NADPH-dependent conversion of 3′-hydroxylated and 4′-*O*-methylated isoflavones into MDB-containing products.

The MDB-forming reaction converting calycosin to pseudobaptigenin (Figure 1) was originally described by Clemens and Barz (18). Microsomal preparations from elicited chickpea (*Cicer arietinum*) cell cultures were shown to form pseudobaptigenin in the presence of O_2_ and NADPH. The reactions were heavily inhibited by characteristic inhibitors of cytochrome P450s, including reversible inhibition by CO, and near-complete inhibition by juglone and plumbagin. The microsomal preparation also catalyzed the parallel MDB formation of 5-hydroxypseudobaptigenin from pratensein. Similar reaction parameters, including kinetics and pH and temperature optima led researchers to conclude that a single cytochrome P450 was responsible for both reactions. However, three decades later, the identity of this proposed cytochrome P450 has remained elusive.

The discovery of cytochrome P450s capable of MDB formation appears to be scattered across the large array of specialized pathways. In the phenylpropanoid-derived lignan pathway, dual-function CYP81Q1 from *Sesamum indicum* catalyzes two sequential MDB formations to form (+)-sesamin, the major lignan in sesame seeds (19). In the tyrosine-derived benzylisoquinoline alkaloid (BIA) pathway, three members of the CYP719A subfamily catalyze MDB formations on (*S*)-scoulerine and its protoberberine derivatives, in the berberine and sanguinarine pathways (20–22). Elsewhere, in bacterial natural product biosynthesis, *Streptomyces spectabilis* StvP2 catalyzes a similar reaction during streptovaricin C biosynthesis, a rifamycin-related antibiotic (23). Indeed, MDB formation, specifically by cytochrome P450s, appears sporadically and in disparate chemical structures.

Notably, pseudobaptigenin synthase (PbS) is thought to catalyze the final unresolved step by forming an MDB in (-)-maackiain biosynthesis. In this study, we integrated metabolomic and transcriptomic datasets from *T. pratense* roots following fungal inoculation by hemibio-and necro-trophic fungi to identify candidate genes. Co-expression and network analyses highlighted a subset of cytochrome P450s which were screened for PbS activity. One gene, CYP76F319 (NCBI gene accession LOC123894475), was functionally characterized as *Pseudobaptigenin synthase* (*TpPbS*) using yeast biotransformation assays analyzed by liquid chromatography/mass spectrometry (LC/MS). The complete resolution of (-)-maackiain biosynthesis will facilitate its *de novo* heterologous production in engineered microbial chassis or other platforms. Furthermore, access to this structure and its derivatives will accelerate the characterization of their bioactivity in health and agriculture.

## RESULTS

We sought to investigate the metabolic shift in *T. pratense* in response to infection by fungal pathogens with different lifestyles. Two-week-old seedlings grown in semi-solid MS media were inoculated with agar plugs of either *Fusarium oxysporum* (hemibiotrophe), *Phoma medicaginis* (necrotrophe), or a mock control. Short (3 dpi) and long (7 dpi) incubations were conducted to span the biotrophic and necrotrophic phases of *F. oxysporum* and the resulting tissues for all treatments were harvested for metabolite and RNA extraction and subsequent analysis.

### Pathogen-specific remodeling of root isoflavonoid profile

Metabolomic analyses identified a diverse set of (iso)flavonoids across the samples, with unique chemical “fingerprints.” We used authentic chemical standards to detect and quantify the isoflavone aglycones (genistein, daidzein, glycitein), their corresponding glucosides (genistin, daidzin, glycitin), the *O*-methylated derivatives (formononetin, biochanin A), and the pterocarpans ((-)-maackiain, trifolirhizin) (Figure 2). Among these, biochanin A and formononetin were the highest accumulating across most treatments, exhibiting baseline levels of 103 ± 69 and 84 ± 49 µg/mg fresh weight (FW), respectively, in the mock samples (Figure S1). The exception to this were PhomaL tissues, which became depleted in the *O*-methylated compounds in favour of glycitin accumulation up to 67 ± 43 µg/mg FW. A few other targets including daidzin, (-)-maackiain, and trifolirhizin, became significantly enriched in the PhomaL treatment (7-dpi), relative to PhomaS (3-dpi) or the mock control.

**Fig. 2.**
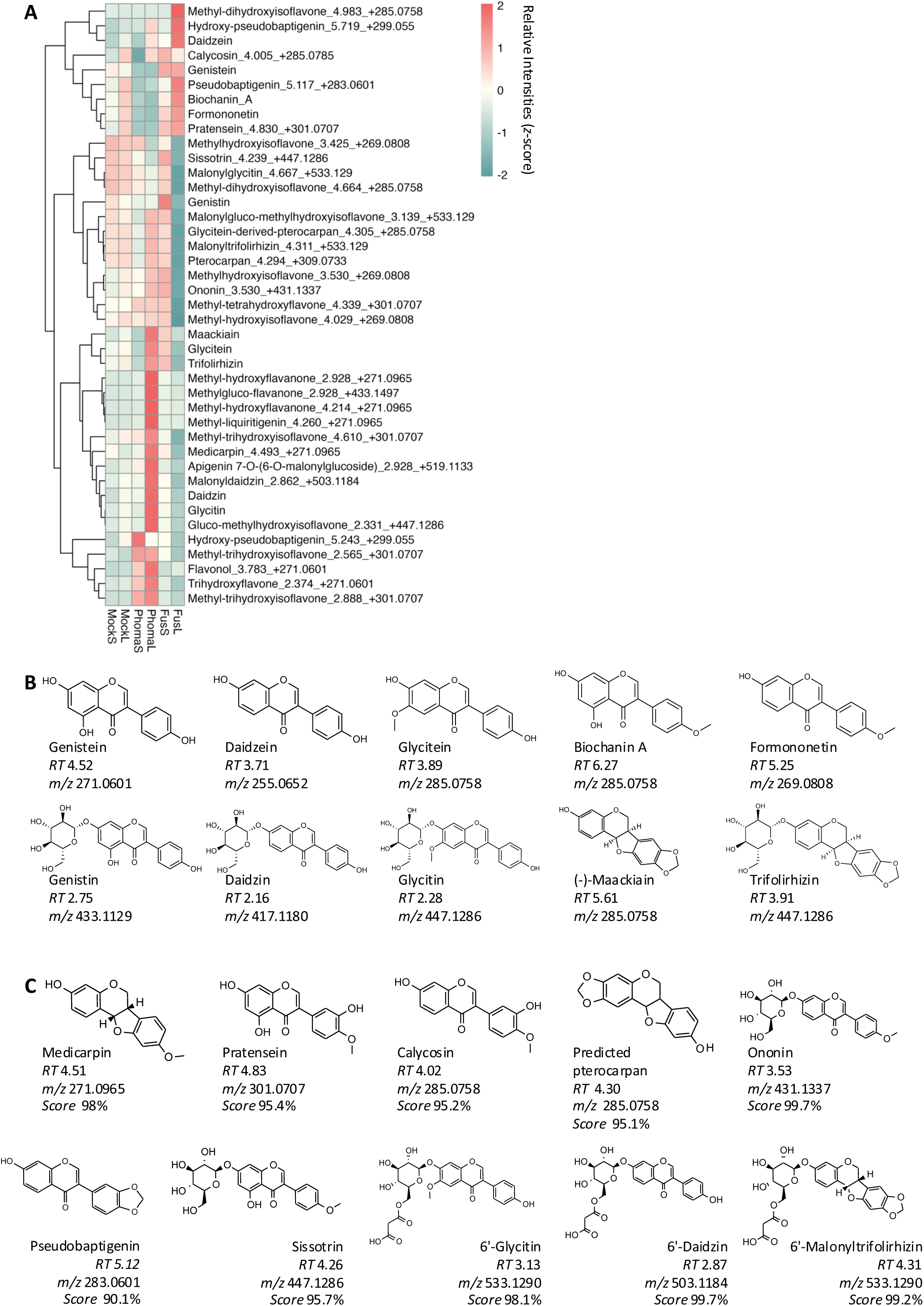
Infection-induced changes in root isoflavonoid composition. **(A)** Heatmap of relative peak intensities (*z-*scores) of known and predicted isoflavonoids normalized across compounds. Labels are annotated with compound names for those verified against authentic standards, or putative identities or subclasses with *rt* and [M+H]^+^ for others. **(B)** Chemical structures of the ten compounds quantified with authentic standards. **(C)** Putatively identified isoflavonoids based on accurate [M+H]^+^ and MS/MS fragmentation spectral matching. A threshold of CSI:FingerID score of >95% was set for identification.

An additional 31 unknown compounds were classified by predicted chemical classes based on their MS/MS fragmentation profiles (Table S1). A subset of 13 predicted features could be putatively identified as calycosin, pratensein, pseudobaptigenin, ononin (glucosyl-formononetin), sissotrin (glucosyl-biochanin A), 6’-malonylglycitin, 6’-malonyldaidzin, and 6’-malonyltrifolirhizin, as well as the flavonoids methyl-liquiritigenin and apigenin 7-*O-*malonylglucoside (Figure S2A, Table S2). The identities of putative calycosin, pratensein, and pseudobaptigenin peaks were subsequently confirmed by matching their exact mass, isotopic pattern, retention time (*rt*), and fragmentation pattern to those of authentic standards (Figure S2B). The pterocarpans, (-)-maackiain, trifolirhizin, and (-)-medicarpin exhibited a similar profile, accumulating to the highest levels in PhomaL roots (Figure S1A). There were three additional predicted pterocarpan derivatives (Table S1), including a putative glycitein-derived pterocarpan, with the same *m/z* as (-)-maackiain, but eluting at a distinct *rt* (Figure 2C, Figure S2A).

### Transcriptional responses to fungal infection

RNA sequencing was performed on the same set of samples using a reference-based transcriptomics approach. Mock samples exhibited high quality mapping to the *T. pratense* reference genome (RefSeq accession: GCF_020283565.1), with an average of 87.6 ± 0.1% of reads uniquely mapping to the genome. In contrast, fungal-infected samples displayed a reduced proportion of uniquely mapped reads (FusS, 61.9 ± 14.2%; FusL, 5.4 ± 2.1%; PhomaS, 64.2 ± 14.4 %; PhomaL, 75.4 ± 9.5%) (Figure S3A). In the case of *Fusarium*-infected samples, the proportion of reads mapping to the *T. pratense* and *F. oxysporum* Fo47 (RefSeq accession: GCF_013085055.1) genomes was inversely related (87.8 ± 2.2%) (Figure S3B). This indicates that the low host (*T. pratense*) mapping rate could be attributed to the presence of fungal RNA in the necrotic tissue, rather than poor RNA quality or sequencing coverage.

Differential gene expression (DESeq2) analysis of treated versus mock samples revealed the greatest transcriptomic shift in response to *Fusarium* infection (Figure 3). The FusL group displayed both the largest total number (5681 up; 7818 down) and the greatest intersection (3511 up; 4952 down) of differentially expressed genes (DEGs). A comparison of FusL relative to FusS revealed a large shift in up-and downregulated genes (4241 up; 5445 down); whereas, *Phoma*-infected samples displayed rather minor differences between harvest days (83 up; 61 down) (Figure S4). Overall, fungal treatment led to activation of central metabolism (glycolysis, gluconeogenesis, and pentose phosphate pathway), as well as amino acid and phenylpropanoid biosynthesis, both of which can feed into isoflavonoid production (Figure 3A). Further, flavonoid metabolism (11 genes) was notably enriched in *Phoma*-infected samples. In contrast, pathways related to cellular maintenance (i.e. DNA replication, ribosomal function, motor proteins, and photosynthesis) were broadly downregulated across all groups except for PhomaS. A gene ontology (GO) term enrichment analysis (Benjamini *p-*Adj<0.05) identified 108 upregulated processes, many of which were associated with stress or defence responses, including fungal defence, chitin response, and ethylene signaling (Figure S5). The 102 downregulated pathways were dominated by photosynthesis-and cell cycle-related processes. Generally, FusS (64 up; 41 down) and FusL (87 up; 93 down) were enriched in the majority of the GO terms detected, while PhomaS (9 up; 4 down) and PhomaL (25 up; 48 down) were enriched in fewer.

**Fig. 3.**
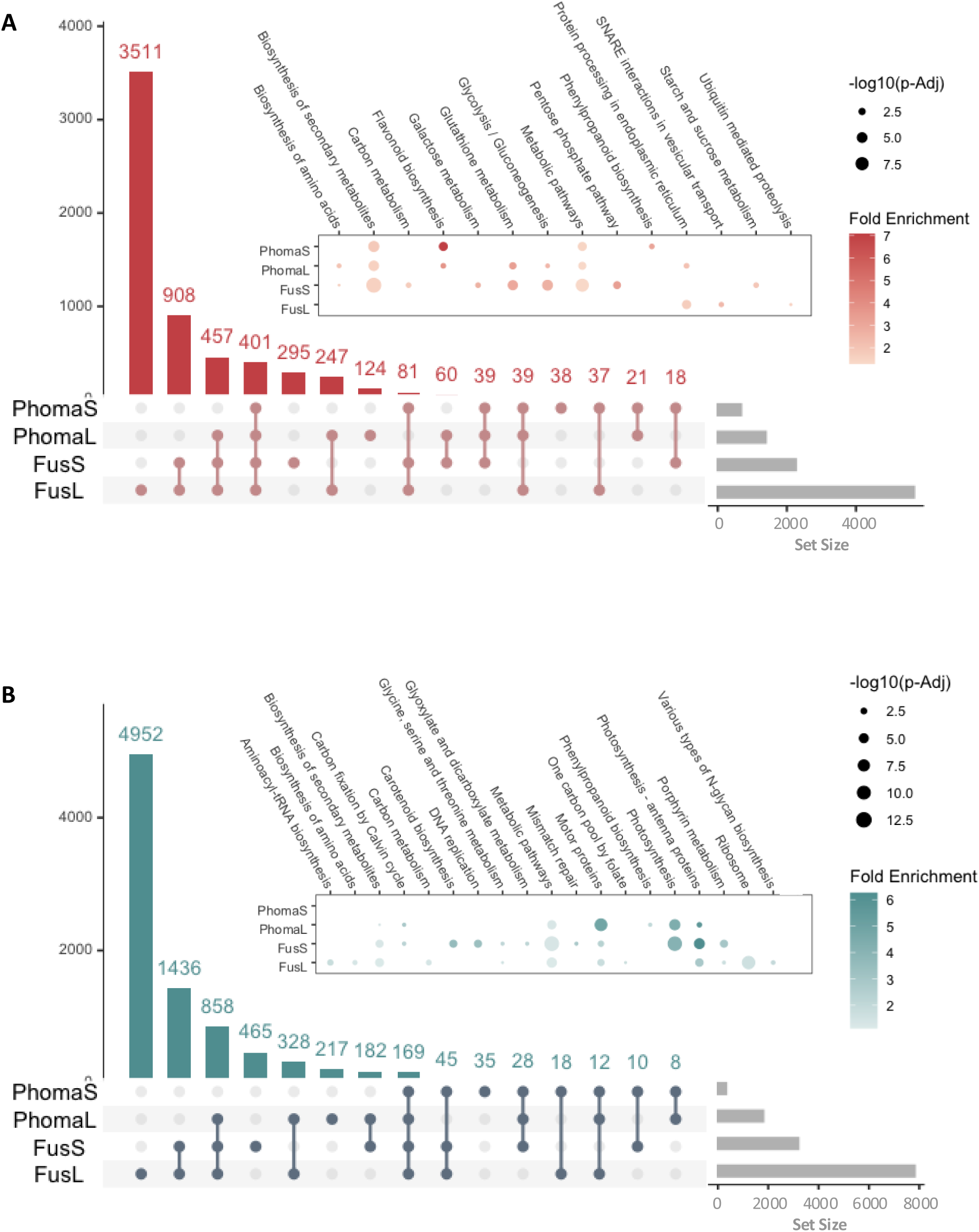
Differential gene expression across fungal-infected roots. UpSet plots and corresponding enriched KEGG pathways (*Benjamini p-*Adj<0.05) of **(A)** upregulated and **(B)** downregulated genes in infected tissues relative to mock roots.

### Differential activation of isoflavonoid pathway genes

A list of 47 *T. pratense* transcripts predicted to encode isoflavonoid biosynthetic genes was compiled based on homology to characterized orthologs, of which 33 were detected as DEGs in at least one group relative to the mock samples (Figure 4). These pathway-associated targets had largely consistent expression trends across most fungal groups, with various pathway DEGs being upregulated in FusS, and PhomaS/L. Further comparison of variance-stabilized expression revealed dynamic regulation of early-, mid-, and late-modules across the five groups (Figure 4A). The early module of the pathway exhibited strong upregulation, including the legume-characteristic chalcone reductase (*CHR*), which, in combination with *chalcone synthase* (*CHS*), produces 6’-deoxychalcone (isoliquiritigenin), the precursor to daidzein. Mid-module transcripts (*IFS, isoflavone 4’-O-methyltransferase* (*HI4’OMT*), and *2-hydroxyisoflavanone dehydratase* (*HIDH*)) displayed similar expression across most treatments, with the exception of *I3’H* which was enriched in FusL. The final module of transcripts (*I2’H, IFR, I4R, PTS* and *isoflav-3-ene synthase* (*I3S*)) was broadly upregulated in FusS and PhomaS/L samples. In contrast, FusL possessed a distinct transcriptional profile, with 13 pathway DEGs being uniquely regulated, the majority (10) of which were significantly downregulated (Figure 4B).

**Fig. 4.**
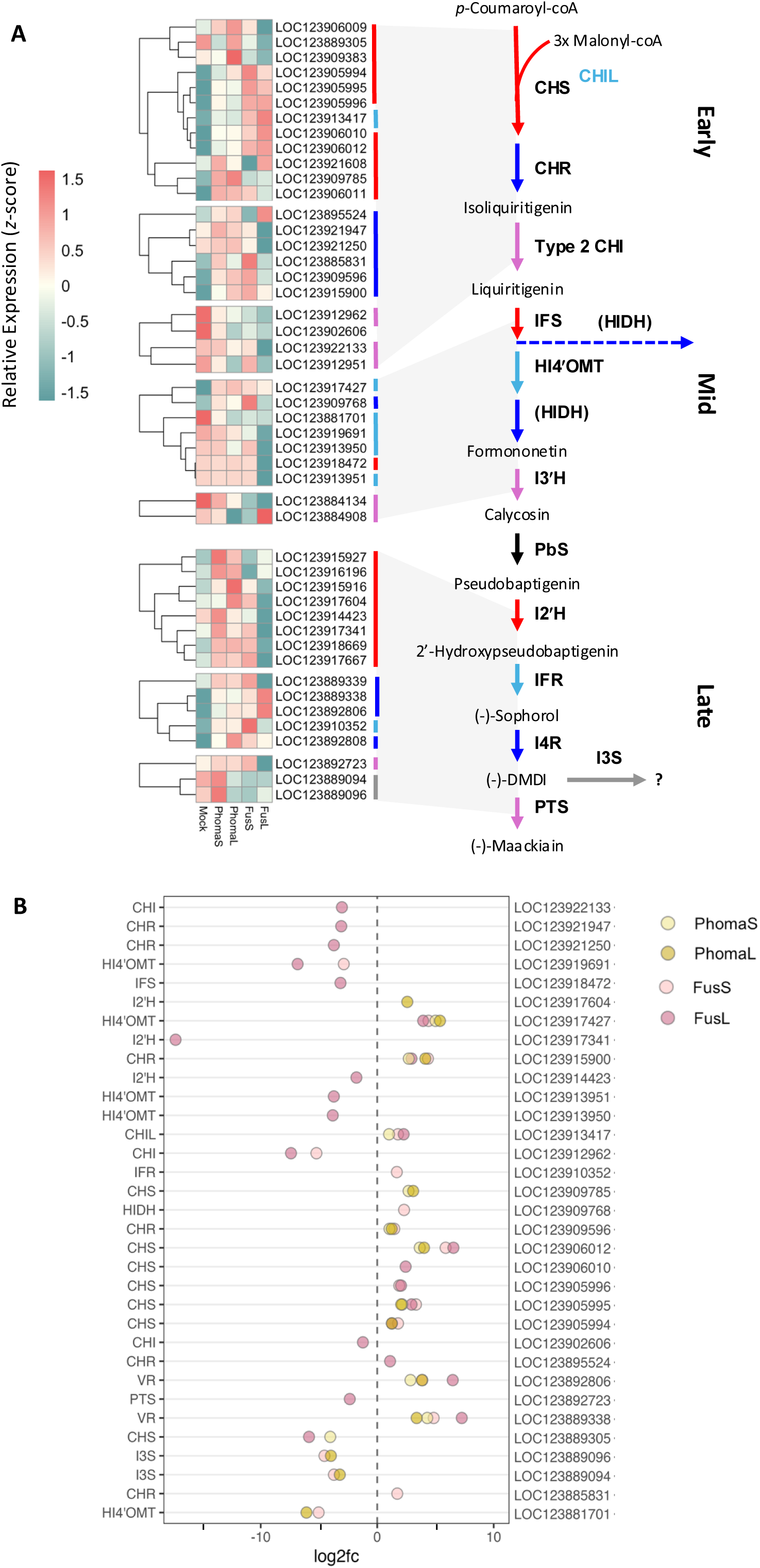
Expression patterns of isoflavonoid pathway-related genes. **(A)** Heatmap of relative variance-stabilized expression (*z*-scores) of isoflavonoid pathway genes, normalized across genes. Early-, mid-, and late-module genes are indicated for the complete pathway from *p*-coumaroyl-coA to (-)-maackiain. **(B)** Log_2_FoldChange (Log2FC) of isoflavonoid pathway DEGs.

### Pseudobaptigenin synthase P450 candidate search

A weighted gene co-expression network analysis (WGCNA) identified clusters of transcripts sharing similar expression profiles to characterized or predicted pathway genes. Hierarchical clustering of samples based on global expression profiles identified FusL samples as outliers in the dataset, and these were therefore excluded from the remaining analyses. The resulting network identified 34 distinct module eigengenes (MEs) (Figure S6), each exhibiting unique patterns of correlation with the different sample groups (Fig. 5A).

**Fig. 5.**
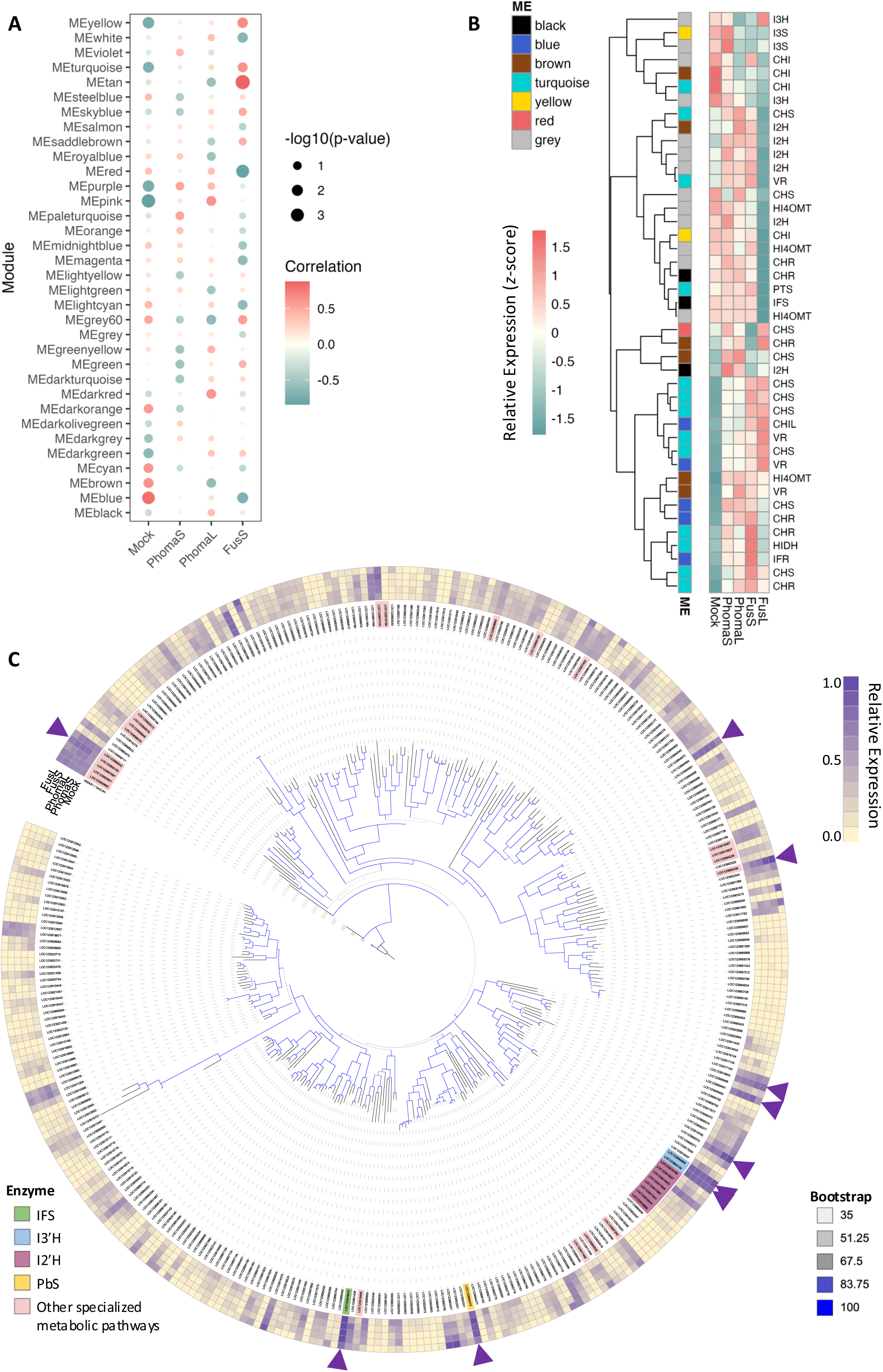
Integrative co-expression and phylogenetic analyses to identify candidate isoflavonoid biosynthetic genes. **(A)** Correlations between the 34 WGCNA module eigengenes (MEs) and each treatment group. FusL samples were excluded from the WGCNA analyses. **(B)** Heatmap of predicted isoflavonoid pathway genes, annotated by ME assignments. **(C)** Maximum-likelihood phylogenetic tree of all 329 P450 genes detected in this study, rooted by the outgroup *Glycine max GmC4H* (NCBI accession: X92437.1). Branches are coloured based on bootstrap confidence, based on 1000 bootstrap replicates. Corresponding variance-stabilized expression normalized within treatment groups is plotted as a heatmap, with the ten most highly expressed P450s across all samples indicated with purple triangles along the perimeter. P450s of interest are annotated: *TpIFS*, green; *TpI3’H*, blue; *TpI3’H*, pink; other P450s annotated as specialized metabolic KEGG pathway genes are indicated in salmon; and *TpPbS* characterized in this study is shown in yellow.

The black and brown modules contained several notable P450-encoding transcripts, including *TpIFS* and a *TpI2’H*, in addition to other pathway genes such as *CHR*, *HI4’OMT* and *I4R* (Figure 5B). Additionally, the turquoise module contained transcripts encoding *I4R* and, critically, *PTS*. The grey module was excluded from further consideration as it represents genes that could not be confidently assigned to any other module. Modules of interest were further filtered for P450 transcripts exhibiting similar correlation values to the ME as the bait genes.

In parallel, variance-stabilized expression of all 329 P450 transcripts detected in this study was overlaid atop a maximum-likelihood phylogenetic tree to visualize the expression landscape of diverse P450s in *T. pratense* roots (Figure 5C). The ten most highly expressed P450s were distributed across the phylogeny, with five genes clustering within a large clade. Two of these clustered genes were predicted *TpI3’H* transcripts, neither of which were detected as DEGs. Similarly, *TpIFS* was not upregulated as a DEG, but ranked as the most abundantly expressed P450 overall (Table S3).

Together, the module-based associations and expression patterns guided the curation and triaging of candidate P450s for subsequent PbS activity screening. Of the five selected candidate genes, two were members of the black ME (P1, LOC123894475; P3, LOC123906377), two were in the brown ME (P2, LOC123909586; P5, LOC123894994), and one belonged to the turquoise ME (P4, LOC123900153). Three candidates, P1, P4, and P5 were highly expressed across all samples, with mean variance-stabilized gene counts 13.7, 12.2, and 12.4, respectively, and were amongst the ten most highly expressed P450s (Table S3). The remaining two candidates exhibited more moderate expression (P3, 8.00; P2, 6.77). All five candidates belonged to different P450 subfamilies (P1, CYP76F; P2, CYP89A; P3, CYP81E; P4, CYP73A; P5; CYP72A), exhibiting <35% sequence identity with each other (Table S4).

### Pseudobaptigenin synthase functional characterization

The five candidate P450 genes were each co-expressed with a compatible plant CPR (*Catharanthus roseus CrCPR*) in yeast (*Saccharomyces cerevisiae*, strain BY4741*-pep4*). Biotransformation assays were conducted with supplementation of 100 µM calycosin for 48 h. Only the strain expressing “P1” produced a new peak corresponding to the expected [281]^-^ *m/z* of pseudobaptigenin (Figure 6A). Furthermore, long (5-day) P1 biotransformation assays supplemented with 150 µM calycosin produced a peak with *rt*, exact *m/z* ([281.0455]^-^), and fragmentation profile identical to the pseudobaptigenin standard (Figure 6B). Therefore, P1 was renamed pseudobaptigenin synthase (TpPbS) and annotated as CYP76F319 (correspondence with Dr. D. Nelson), based on homology to the CYP76F subfamily (Table S5) (24). We further assayed TpPbS with 150 µM of pratensein (5-hydroxycalycosin), which resulted in a new peak with *m*/*z* ([297]^-^) corresponding to 5-hydroxypseudobaptigenin (Figure 6C). We cannot rule out a potential catalytic activity for P2 and P4, as they had limited to no detection in immunoblots of the respective microsomal preparations (Figure 6D).

**Fig. 6.**
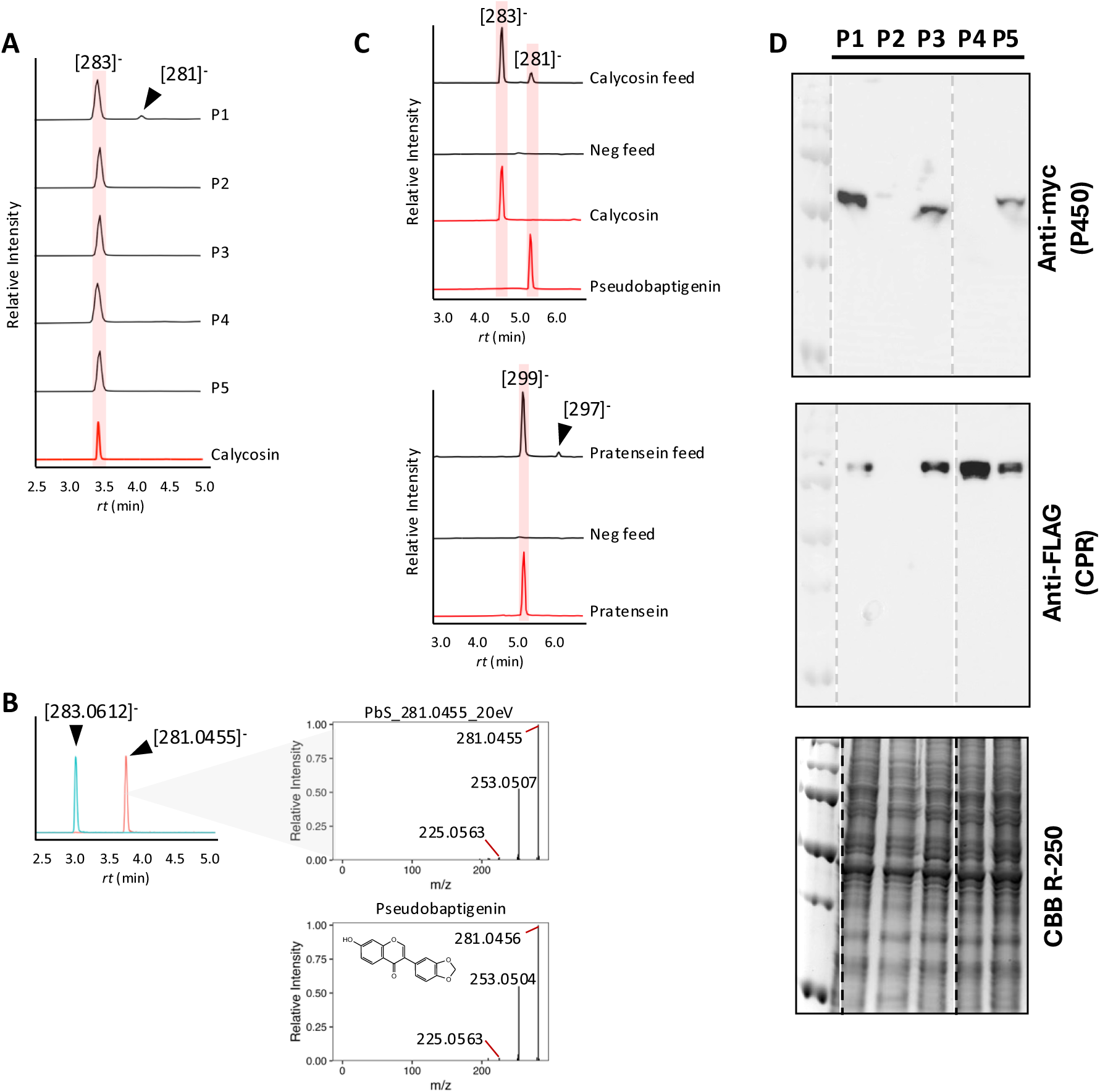
Characterizing a P450 with pseudobaptigenin synthase (PbS) activity. **(A)** LC-TQ/MS single ion monitoring (SIM; [283]^-^ and [281]^-^)-analysis extracts of yeast expressing candidate P450s fed with 100 µM calycosin for 48 h. **(B)** The P1-expressing strain was further biotransformed with 150 µM substrate for 5 d, yielding a product with an exact *m/z* and MS/MS spectral match (CE: 20 V) to the expected [281.0455]^-^ of pseudobaptigenin on LC-QTOF/MS. **(C)** The substrate specificity of P1 was tested with 150 µM of the related substrates, calycosin and pratensein. Extracts were analyzed by LC-TQ/MS in SIM mode for putative peaks corresponding either to calycosin [283]^-^ and pseudobaptigenin [281]^-^, or pratensein [299]^-^ and 5-hydroxypseudobaptigenin [297]^-^. **(D)** P450 expression was validated by SDS-PAGE and subsequent immunoblots (anti-FLAG or anti-myc) or total protein stain (CBB R-250) of microsomal preparations from 24 h induced cultures.

## DISCUSSION

(-)-Maackiain straddles the boundary between phytoalexin and “phytoanticipin,” the latter term was coined by J.W. Mansfield to encompass low molecular weight, antimicrobial compounds that are preformed, prior to infection, or produced directly thereafter from preexisting constituents (25). Indeed, this definition allows (-)-maackiain to be considered preformed, as it exists in the roots of *T. pratense* in the glucoside-conjugated form and is “released” by a glucosidase upon tissue maceration and decompartmentalization (11, 26). Conversely, (-)-maackiain is also synthesized *de novo* in *T. pratense* in response to microbial pathogens or molecular elicitors (27, 28). In our study, this duality explains the constitutive expression of many pathway enzymes, and even entire modules, while there is an apparent elicited increase in flux through to (-)-maackiain and other defence-related pterocarpans in some infected tissues.

In previous studies, the expression of pterocarpan biosynthetic genes have been shown to be induced by elicitor treatment, followed by an accumulation of cognate metabolites. For example, soybean pterocarpan synthase (*GmPTS1*) and downstream glyceollin biosynthetic genes were induced by yeast-extract (YE) treatment of soybean cell-suspension cultures (13, 29). However, *GmPTS1* transcript levels were reduced after their peak at 10 h post-treatment, followed by the accumulation of glyceollins (13). Such rapid transcript profile shifts might be missed in the longer incubation times we have used, although fungal colonization of roots via agar plug inoculation is expected to progress more slowly than elicitation in cell suspension systems.

Treatment with the fungal pathogens, *F. oxysporum* and *P. medicaginis*, did not lead to a cohort of tightly co-upregulated biosynthetic genes in the isoflavone-to-pterocarpan module, consistent with the complex and branched regulation of (-)-maackiain biosynthesis described above. Accordingly, homologs of known pathway genes were not uniformly upregulated across treatments. Nonetheless, an analysis of normalized gene counts indicated an enrichment for the late-module genes in *Phoma*-infected and FusS tissues (Figure 4A). This transcriptional activation was aligned closely with the pronounced accumulation of end-state metabolites, (-)-maackiain and trifolirhizin, particularly in PhomaL (Figure 2A). In contrast, late-stage Fusarium infection (FusL) triggered a broad downregulation of isoflavonoid pathway genes, correlating with a marked bottleneck at early intermediates such as daidzein and formononetin. In this manner, different fungal pathogens elicited distinct metabolic profiles that reflect pathogen-specific reprogramming of isoflavonoid metabolism, defence responses, and cellular homeostasis.

A WGCNA analysis assigned bait enzymes into co-expression modules (black, brown, and turquoise) that included characterized biosynthetic genes and candidate P450s. The curated list was further filtered and supplemented by analyzing normalized read counts, phylogeny, and binary comparisons between treatments. In particular, *TpIFS* was found in the “black” module and was also the most strongly expressed P450 across the whole dataset (Figure 5B; Table S3). Similarly in chickpea, the orthologous *IFS* was highly expressed in a resistant cultivar (30). Other members of this module included a predicted *TpI2’H* (LOC123915927) and our newly identified *TpPbS.* The list of most highly expressed P450 transcripts further contained two putative *TpI3’H* transcripts, indicating robust activation of pterocarpan biosynthesis in *T. pratense* roots grown in media.

The *T. pratense* PbS was classified as CYP76F319, despite all previously characterized MDB-forming plant enzymes belonging to the CYP81 or CYP719 families, to our knowledge (Table S5). The CYP76 family was originally identified in eggplant seedlings (CYP76A) (31), and known activities range from dealkylation of xenobiotics to hydroxylation or oxidization of terpenoid intermediates (32–34). This functional diversity is mirrored in the size of the CYP76 family, which currently comprises 55 subfamilies, of which CYP76F is the largest (24). This subfamily features catalysts in terpenoid and polyphenol biosynthesis, including CYP76F112 (*Ficus carica*), which catalyzes a distinctive 5-membered-ring cyclization converting demethylmarmesin into marmesin (35–38).

The mechanism of P450-catalyzed MDB formations is not well understood. Homology modelling and mutagenesis of *S. indicum* CYP81Q1 indicated Ala/Ser308 as a crucial residue, differing from the canonical Thr308, stabilizing water molecules participating in hydrogen-bond networks (39). This network is thought to orient and possibly activate the substrate for an intramolecular nucleophilic attack between proximal hydroxyl and methoxy groups, leading to MDB formation. Despite the presence of the same MDB-containing moieties in their respective substrates (1,3-benzodioxol), TpPbS/CYP76F319 and CYP81Q1 appear to employ different mechanisms, as the former possesses the canonical Thr308. However, multiple MDB-forming routes must be available, as the bacterial SsStvP2 acts on proximal methoxy and keto groups. Based on the crystal structure of the holoenzyme, SsStvP2 is proposed to activate the methoxy group via a water-mediated proton relay involving a catalytic triad, which in turn triggers tautomerization and intramolecular nucleophilic substitution, resulting in MDB formation (23).

Our characterization of CYP76F319 closely aligns with the early description of the PbS-catalyzed reaction in yeast-elicited chickpea cell cultures (18). Corroborating their work, we found that TpPbS accepts both calycosin and pratensein (5-hydroxy-calycosin) (Figure 6C).

Indeed, the discovery of (-)-maackiain and its biosynthetic route has greatly benefited from fungal elicitation and plant-derived cell culture. As early as 1976, J.L. Ingham described the accumulation of “induced” isoflavonoids, (-)-maackiain and (-)-medicarpin, in response to fungal inoculation of chickpea stems (6). This was followed by compound isolation using fungal-elicited chickpea cell cultures (40) and, subsequently, pathway characterization in *T. pratense* and chickpea (41–43). Callus cultures derived from *Maackia amurensis* have also been shown to accumulate high levels of (-)-maackiain and (-)-medicarpin, along with retusin, daidzein, genistein, and formononetin (44). The inflorescence-derived callus accumulated 5.8 mg/g (DW) of (-)-maackiain, which was 17-fold greater than concentrations found in *M. amurensis* heartwood, where the compound was originally discovered (44–46). These results suggest that cell culture or heterologous platforms could viably permit large-scale production of these chemistries.

The two fungal species chosen for this study are well-characterized for their pathogenicity on *T. pratense*, with our test specimens having been extracted nearly five decades ago from the roots of the species and maintained by the Canadian Collection of Fungal Cultures. The necrotrophe, *Phoma medicaginis* affects many food and forage legumes, e.g., causing “spring black stem” and “leaf spot” disease in alfalfa (*Medicago sativa*), and frequently co-occurs with or precedes pea aphid (*Acyrthosiphon pisum*) infestation (47, 48). Upon *P. medicaginis* exposure, resistant varieties of alfalfa upregulate an array of genes associated with specialized metabolism, particularly phenylalanine ammonia lyase (PAL) and IFR (49). Other studies have shown a robust metabolite response in alfalfa, including the accumulation of pterocarpans like (-)-medicarpin, which inhibits *P. medicaginis* mycelial growth (50, 51). Our data indicate a similar response in *T. pratense* roots, which became enriched with (-)-medicarpin, (-)-maackiain, and other (iso)flavonoids upon *P. medicaginis* infection (Figure 2).

Similarly, pathogenic members of the *Fusarium* genus are responsible for vascular wilt disease in many important crops, including legumes. In other studies, a concurrent reduction in trifolirhizin and an increase in (-)-maackiain has been attributed to the liberation of the latter from the glycoside (11), contributing to inhibition of germ tube and radial mycelia growth in *F. roseum* (52). Here, the hemibiotrophic infection strategy employed by *F. oxysporum* leads to a sequential cascade of metabolic shifts in the roots. The (iso)flavonoid profile at 3 dpi (FusS) was most similar to the mock, whereas most target metabolites were drastically reduced at 7 dpi (FusL) (Figure 2), suggesting a collapse of the plant defences as necrosis advances. The differences in isoflavonoid metabolic responses to necrotrophy and hemibiotrophy might stem from the ability to rapidly perceive a hostile pathogen and mount chemical defences, before fungal counter-measures take effect.

Some pathogenic fungi, including *Fusarium* species, have developed detoxification strategies that modify and neutralize these plant antimicrobial metabolites. For instance, the pterocarpans, (-)-maackiain and (-)-medicarpin, can be hydroxylated and/or demethylated by pathogens, including *B. cinerea, Fusarium* spp., and *Colletotrichum* spp., resulting in reduced bioactivity (53–56). A few pathogens (e.g., *Nectria haematococca*, the anamorph of *F. solani*) have been noted for their ability to additionally catalyze C-ring fissions of (-)-maackiain and (-)-medicarpin to produce the isoflavanones sophorol and vestitone, respectively (54). Such conversions represent the reverse reactions carried out by the plant biosynthetic enzymes I4R and PTS. More specifically, *N. haematococca* genes have been linked to its ability to metabolize (-)-maackiain (*MAK1, MAK2,* and *MAK3*), with *MAK1* having been cloned and annotated as a flavin-containing monooxygenase (57, 58). Such detoxification mechanisms may, therefore, contribute to the differences in phytoalexin profiles observed at 3 and 7 dpi.

Following the elucidation of the final unknown PbS step of (-)-maackiain biosynthesis, our integrated multi-omics datasets can serve as a valuable resource for future gene mining efforts targeting additional components of pterocarpan metabolism. Multi-omics approaches have rapidly enabled the discovery of biosynthetic enzymes across specialized metabolism from isoflavonoids to alkaloids, terpenoids, and beyond (59–63). Indeed, a flurry of recent publications reported the final reactions leading to the pterocarpan subclass of (-)-glyceollins (13, 64–66). Collectively, seven glyceollin synthase cytochrome P450s were identified, capable of generating at least four new scaffolds from two substrates. Conversely, there have been a limited number of *T. pratense* transcriptomic studies, with even fewer leveraging these resources to describe the biosynthesis and regulation of its unique isoflavonoid profile (67–70). Here, we present a dataset of pathogen-induced *T. pratense* roots and demonstrate its utility in uncovering a key pathway enzyme. The discovery of TpPbS/CYP76F319 enables reconstruction of the complete (-)-maackiain pathway for the first time, paving the way to optimized bioproduction in microbial or plant chassis. Finding access to (-)-maackiain and its derivatives, including trifolirhizin, will open a new portfolio of antimicrobial compounds for health and agricultural applications.

## MATERIALS AND METHODS

### Chemicals

Analytical and assay grade standards were purchased for biotransformation assays, including calycosin (≥98□%, Cayman Chemical, MI, USA, Cat. 20929), pseudobaptigenin (98.41%, MedChemExpress, NJ, USA, Cat. HY-N10616), pratensein (99.92%, MedChemExpress, Cat. HY-N7981), and fluorescein sodium salt (≥97.5□%, Sigma-Aldrich, Cat. 30181). The following chemicals were used as authentic chemical standards for peak identification: genistein (≥99□%, Thermo Fisher Scientific, Cat. J63241.MC), genistin (≥95%, Sigma-Aldrich, Cat. G0897), daidzein (≥97□%, Thermo Fisher Scientific, Cat. B22877), daidzin (≥98%, Sigma-Aldrich, Cat. 42926), glycitein (≥97%, Sigma-Aldrich, Cat. G2785), biochanin A (≥95%, Sigma-Aldrich, Cat. PHL80012), formononetin (≥98%, Sigma-Aldrich, Cat. 94334), (-)-maackiain (≥98%, Toronto Research Chemicals, Cat. TRC-M214390), and trifolirhizin (≥98□%, Cayman Chemical, Cat. 34655). Metabolites were stored at −20 °C in lyophilized form or dissolved in DMSO.

### Fungal material and growth

The fungal specimens were received from the Canadian Collection of Fungal Cultures (DAOMC) kept by Agriculture and Agri-Food Canada (AAFC). The species included *F. oxysporum* (DAOMC-174256; CCFC003211) and *P. medicaginis* (DAOMC-174525; CCFC003226). They were both initially isolated from *T. pratense* roots growing in Ste-Anne-de-Bellevue, Québec, Canada, by P. Tettch, on the farms of Macdonald College (now the Macdonald Campus of McGill University) in 1979. Both *F. oxysporum* and *P. medicaginis* (syn. *Ascochyta medicaginicola*) are ascomycetes are containment level 1 (CL1) species. Fungi were cultured on potato dextrose solid agar (PDA) media at 25 °C, with a 16:8 h light:dark cycle and 50 % humidity for two weeks prior to plant inoculation.

### Plant material and inoculation

*Trifolium pratense* (cv. Start) seeds were bleach-sterilized and germinated in the dark for 10 d, prior to their transfer into sterile semi-solid, 1/2x Murashige and Skoog (MS) media (2.2 g/L MS salts with Gamborg’s vitamins (Sigma-Aldrich, Cat. M0404), 20 g/L sucrose, and 0.8 g/L Phytagel in magenta boxes. Seedlings were grown in a tissue culture chamber (22 °C; 16:8 h light:dark cycle; 50 % humidity) for 3 weeks. A single PDA plug (1 cm x 1 cm) taken from the growing fungal hyphal edges was used to inoculate the media at the base of each stem, while agar plugs of pure PDA were used for mock controls. Root tissues were harvested at 3 and 7 dpi (minimum *n*=4 for each condition), and homogenized using mortar and pestle with liquid N_2_, before being divided for subsequent metabolite and RNA extraction and stored at <u>−</u>80 °C.

### Metabolite extraction and analysis

#### Metabolite extraction

Frozen pre-homogenized root tissues were thawed and extracted in 20 µL/mg fresh weight (FW) of 80% MeOH. Samples were vortexed and sonicated in an ultrasonic bath with ice for 20 min, then microcentrifuged at 11,000 x *g* for 10 min to remove cell debris. The supernatants were transferred, dried down, and stored at −20 °C. Prior to LC-QTOF/MS analysis, dried samples were resuspended in 80 % MeOH containing 20 ng/mL fluorescein and filtered through 0.22 µm hydrophobic PTFE syringe filters (Avantor; QC, Canada).

Yeast biotransformation assays were harvested at 2,500 x *g*, cells were washed with LTE buffer (20□mM Tris-HCl (pH 7.5), 200 mM LiOAc, and 2 mM EDTA), then both media and cells were extracted in 75 % EtOH. Cell debris was removed by centrifugation at 21,000 x *g* for 30 min, and clarified supernatants were dried down and then stored at −20 °C. For LC/MS, samples were resuspended in 80 % MeOH containing 20 ng/mL fluorescein as an internal standard.

#### LC-quadrupole-time-of-flight/MS

Both targeted (authentic standards) and untargeted (unknown molecular features) analyses of *T. pratense* metabolite extracts were performed using a 6545 LC-QTOF/MS (Agilent Technologies). First, 1 µL of samples were injected and analytes were chromatographically resolved using a ZORBAX RRHD Eclipse Plus C18 column (2.1 × 50 mm, 1.8 μm) and guard column Agilent ESC18 (2.1 x 50mm, 1.8 μm). The binary solvent system consisted of solvent A (H_2_O + 0.1 % formic acid (FA)) and solvent B (acetonitrile (ACN) + 0.1 % FA), using a flow rate of 0.4 mL/min. The solvent gradient was developed as follows: 10 % B, 0–0.5 min; 50 % B, 6.0 min; 90 % B, 9.10–12.5 min, 100 % B. Samples were run in both positive and negative polarity scan modes with the following parameters: capillary voltage 4,000 V, fragmentor voltage 150 V; skimmer 65 V; sheath gas temperature 250 °C; sheath gas flow 10 L/min; nebulizer 30 psig; scan rate 2 spectras/sec; and a mass range of 100–1,100 *m/z*. Targeted MS/MS fragmentation of additional features was performed using collision energies (CE) 10, 20, and 40 V.

An Agilent Revident LC-QTOF/MS (Agilent Technologies) was used to analyze exact mass and MS/MS fragmentation profiles of products from yeast expressing P1 fed with calycosin, as well as authentic standards (calycosin, pseudobaptigenin, and pratensein). Chromatographic resolution was carried out on an HPLC Poroshell 120 EC-C18 (3.0 ×50 mm, 2.7 M) (Agilent Technologies) with a guard column (Poroshell 120, UHPLC Guard.EC-C18, 3.0 mm) at 30 °C.

A binary solvent system comprised of solvent A (H_2_O + 0.1 % formic acid (FA)) and solvent B (acetonitrile (ACN) + 0.1 % FA), were used at a flow rate of 0.4 mL/min. Sample volumes 1 or 5 µL were injected, before the solvent gradient was developed as follows: 10–30 % B, 0–1 min; 10–50 %, 1–4 min; 50–90 %, 4–5 min; 90–100 %, 5–5.5 min. All other parameters were the same as outlined above.

Tandem MS spectra were analyzed by SIRIUS 4 (71–76), with default parameters. A mass accuracy of 10 ppm was applied for all possible ionizations, using ZODIAC for molecular formula validation, all fallback adducts, and all available databases for structure searches. CANOPUS was employed to predict chemical classes from fragmentation patterns and molecular fingerprints. Putative compound identities were reported for molecular features with a CSI:FingerID score ≥95% or for those that matched the MS/MS spectra of authentic standards, otherwise, only the predicted chemical class was reported.

#### LC-triple-quadrupole/MS

An Agilent 6495 triple quadrupole LC/MS system (Agilent Technologies) was used for initial yeast screening, while an Agilent 6470 triple quadrupole LC/MS system (Agilent Technologies) was used for subsequent analyses of P1 biotransformation assays. All chromatography was carried out on an HPLC Poroshell 120 EC-C18 (3.0 ×50 mm, 2.7□M) (Agilent Technologies) with a guard column (Poroshell 120, UHPLC Guard.EC-C18, 3.0□mm) at 30 °C. The binary solvent system comprised of solvent A (H_2_O + 0.1□% formic acid (FA)) and solvent B (acetonitrile (ACN) +□0.1□% FA), were used at a flow rate of 0.4 mL/min. Sample volumes 1 µL were injected, before the solvent gradient was developed: 10–30□% B, 0–1□min; 10–50□%, 1–4□min; 50–90□%, 4–5□min; 90–100□%, 5–5.5□min. Compounds were ionized by electrospray ionization, and acquisition parameters were set to detect in negative polarity in single ion monitoring (SIM) mode for [281]^-^, [283]^-^, [297]^-^, and [299]^-^; capillary voltage 3,000 V; fragmentor voltage 166 V; sheath gas temperature 250 °C; sheath gas flow 11 L/min; nebulizer 35 psi; and a mass range of 100–1,100 *m/z*.

### RNA isolation, transcriptome sequencing, and gene expression analysis

Total RNA was extracted from homogenized root tissue of each mock-or fungal-infected sample, using the RNeasy Plant Mini Kit (QIAgen, Hilden, Germany, Cat. 74904) according to the manufacturer’s protocol. Total RNA was quantified using a NanoPhotometer® NP80 (Implen, CA, USA), and integrity was confirmed through gel electrophoresis, with 0.2□g of RNA run on a 1□% (w/v) agarose gel with 1□% (v/v) bleach. Extracted RNA (500 ng) was sequenced by Genome Quebec using Illumina NovaSeq 6000 paired-end sequencing.

Trimmed, filtered, and high-quality reads were aligned to the NCBI *T. pratense* (RefSeq accession: GCF_020283565.1) or *F. oxysporum* Fo47 (RefSeq accession: GCF_013085055.1) reference genomes using STAR 2.7.11a. Functional annotations provided with the *T. pratense* reference genome (RefSeq GTF file) were used for gene-level quantification. Differential gene expression was analyzed by DESeq2 for all fungal-infected samples relative to the mock control, and filtered for *p-*Adj<0.05 and an absolute log_2_FoldChange (Log2FC)□>1 to be considered significantly differentially expressed. KEGG pathway and GO term enrichment analyses were conducted using the Database for Annotation, Visualization, and Integrated Discovery (DAVID) bioinformatics tool (NIH), with the threshold for enrichment significance set to *p*-Adj<0.05.

A weighted gene co-expression network analysis (WGCNA; R package) was performed using variance-stabilized gene expression values across all conditions, excluding FusL samples. Genes with low transformed expression (<10) in at least 25 % of samples were filtered out prior to network construction. A soft-thresholding power was selected to approximate scale-free topology, and a network was constructed using a signed topological overlap matrix. Modules with highly correlated (>0.7) eigengenes were merged. Correlations between module eigengenes (MEs) and treatment groups were calculated, and the module memberships of bait pathway genes were examined to identify modules of interest.

All putative *T. pratense* P450 genes were compiled using the HMMER P450 database, then MUSCLE aligned using super5 mode. The resulting multiple sequence alignment was used to construct a maximum-likelihood tree in IQTREE using 1000 bootstrap replicates, using *Glycine max GmC4H* as the outgroup (NCBI accession: X92437.1). Variance-stabilization transformed gene counts were mapped onto the tree and further scaled by treatment group. Genes lacking gene count data were removed from the tree. The transformed expression data was also used to identify the top ten most highly expressed P450s across all samples, including the FusL group.

### Functional characterization of cytochrome P450s

Candidate genes were codon-optimized (Supplementary Materials) for *Saccharomyces cerevisiae* expression (Twist Biosciences, CA, USA) and cloned into the *GAL1* multiple cloning site, in-frame with a C-terminal c-myc tag, in *pESC-Leu2d-GAL10:CrCPR-FLAG*. Constructs were transformed into *S. cerevisiae* BY4741*-pep4* (*MATa his3*Δ*1 leu2*Δ*0 met15*Δ*0 ura3*Δ *pep4*Δ) strain. Biotransformation reactions were conducted as previously described (77). Briefly, overnight cultures were grown in 2 mL synthetic complete (SC) dropout media (2% glucose and appropriate dropout supplements) lacking leucine (SC-Leu) at 30 °C, 225 rpm. These were subcultured into SC-Leu media containing 2□% galactose as the sole carbon source (appropriate dropout supplements) to a final OD of 0.3, and 500 µL of induced cultures were fed 100 µM calycosin in 96-deepwell microtitre plates. Biotransformation assays were incubated for 48 h at 30°C, with shaking at 900 rpm. Further characterization of the P1-expressing strain was performed with some modifications; cultures were induced at a final OD of 3.0, and fed with 150 µM of either calycosin or pratensein, and incubated for 5 d.

### Microsome preparation and immunoblotting

To assess expression of candidate P450s in *S. cerevisiae*, microsomal fractionation was performed (77). Large-scale 50 mL cultures were grown in SC-Leu (2% glucose), then transferred into fresh SC-Leu containing 2% galactose only, for further incubation at 30 °C for 24 h, with shaking at 225 rpm. Harvested cells were resuspended (1:3; w/v) in microsomal extraction buffer (25□mM HEPES (pH 6.8) containing 150□mM LiOAc, 250□mM sucrose, 1□mM phenylmethylsulfonyl fluoride (PMSF), and 1 mM dithiothreitol (DTT)). Cells were mechanically lysed by vortexing with glass beads for 15 ×□30 sec on ice. Lysates were clarified at 10,000 x *g* for 2□min at 4 °C to remove cell debris, then the supernatant was microcentrifuged at 21,000 x *g* for 2□h to pellet the crude microsome fraction. Fractions were resuspended in microsomal extraction buffer, then boiled in sodium dodecyl sulphate (SDS) sample buffer prior to gel electrophoresis.

SDS-polyacrylamide gel electrophoresis (SDS-PAGE) gels were composed of 4□% (w/v) acrylamide stacking and 10□% resolving gels, and electrophoresis was run at 200□V for at least 45□min or until the dye front ran off. Gels were either stained for total protein using Coomassie Brilliant Blue R-250 (CBB R-250) or transferred onto poly(vinylidene) difluoride (PVDF) membrane for immunoblotting. Membranes were blocked in 5□% (w/v) skim milk powder and incubated with mouse anti-FLAG or anti-myc (Sigma Aldrich) diluted 1:3000 (v/v) in tris-buffered saline (pH 7.5) containing 0.1□% Tween (TBST), 1□% (w/v) BSA, and 0.05□% (w/v) NaN_3_. Immunoreactive bands probed with secondary goat anti-mouse antibody conjugated to horseradish peroxidase were treated with Clarity ECL substrates (Bio-Rad, CA, USA) and chemiluminescence emissions were detected by the ChemiDoc imager (Bio-Rad).

## STATISTICS AND REPRODUCIBILITY

All experiments were conducted on a minimum of *n*□=□3 or *n* = 4 biologically independent samples for transcriptomics and metabolomics, respectively. Data is represented as sample mean ± standard deviation. To compare significant differences between all groups, the data were analyzed using one-way analysis of variance (ANOVA) with subsequent pairwise comparisons conducted using Tukey’s HSD test. Statistical significance was determined as *p<*0.05 and is denoted by letters.

## ACKNOWLEDGEMENTS

This work was funded by the Natural Science and Engineering Research Council of Canada (NSERC) Discovery Grant fund to MD (NSERC RGPIN-2021-02817). LC-QTOF/MS analysis on *T. pratense* samples was conducted by SB lab, supported by the Canada Foundation for Innovation - John R. Evans Leaders Fund (Grant # 35318 to SB). We thank Dr. Philippe Seguin for providing *T. pratense* (cv. START) seeds. We thank Tara L. Rintoul at Agriculture and Agri-Food Canada (AAFC) and the Canadian Collection of Fungal Cultures (DAOMC) for providing the fungal specimens (*F. oxysporum* and *P. medicaginis*). Access to an LC-QTOF/MS to analyze yeast-based biotransformation assays was provided by Jean-Francois Roy at the Agilent Center for Excellence in Mississauga, ON. Initial LC-TQ/MS was conducted at McGill-Agilent Partnership Lab (MAPL) and continued through the TRACES Centre (University of Toronto, Scarborough Campus).

## AUTHOR CONTRIBUTIONS

LMR wrote the first draft of the manuscript and designed all experiments. LL performed LC-QTOF runs and LMR analyzed resulting metabolomics data. LMR conducted all other experiments and analyses and generated the figures. SB reviewed the manuscript. MD conceived the study, secured funding, supervised the research, revised the manuscript, and approved it for submission.

## SUPPLEMENTARY INFORMATION

**Supplementary Fig. 1. Quantified isoflavonoids in *T. pratense* root extracts. (A)** Concentrations of compounds quantified against authentic standards. Letters denote significant differences calculated as determined by ANOVA followed by Tukey’s HSD test. **(B)** Heatmap of relative concentrations (*z*-scores) normalized within samples.

**Supplementary Fig. 2. MS/MS fragmentation spectra of putatively identified compounds in *T. pratense*. (A)** The identities of 12 molecular features were predicted based on their MS/MS spectra. Plots are labelled with *rt*, [M+H]^+^, collision energy, and putative chemical structure. **(B)** MS/MS spectra of authentic standards (calycosin, pratensein, and pseudobaptigenin) confirm the identities of three putatively identified compounds. All plots are annotated with the *m/z* values of the three most intense peaks.

**Supplementary Fig. 3. RNA-Seq STAR alignment statistics.** Visualizing the proportion of multi-mapped, uniquely mapped, and unmapped reads to the **(A)** *T. pratense* NCBI reference genome (RefSeq accession: GCF_020283565.1) and the **(B)** *F. oxysporum* Fo47 reference genome (RefSeq accession: GCF_013085055.1).

**Supplementary Fig. 4. Transcriptional reprogramming in fungal-infected *T. pratense* roots. (A)** DEG volcano plots of infected tissues relative to mock, or long (7 dpi) relative to short (3 dpi) infection time tissues. Significant genes (*p*<0.05) are annotated in red. **(B)** PCA plot of the 15 samples constructed using variance-stabilized gene expression data.

**Supplementary Fig. 5. GO term functional enrichment in response to infection. (A)** Enriched GO terms across up-(red) and downregulated (blue) DEGs (*Benjamini p*-Adj<0.05).

**Supplementary Fig. 6. WGCNA cluster dendrogram.**

**Supplementary Table 1. Classification of 31 molecular features identified in *T. pratense* root extracts.** Compound classes determined based on MS/MS fragmentation matching against natural product fingerprints.

**Supplementary Table 2. Putatively identified compounds.** For each molecular feature, the their corresponding *rt*, [M+H]^+^, and classifications are shown. Features were assigned putative identities when CSI:FingerID scores exceeded 95%.

**Supplementary Table 3. Most highly expressed P450s across all root samples.** Mean and variance were calculated using variance-stabilized gene expression data across all samples.

**Supplementary Table 4. Percent identity matrix of five selected candidate P450s protein sequences.** Generated using ClustalW.

**Supplementary Table 5. Percent identity matrix of the TpPbS protein sequence compared to other P450s of interest.** Generated using ClustalW.

