## Supplementary Information for "Discovery of pseudobaptigenin synthase, completing the (-)-maackiain biosynthetic pathway"

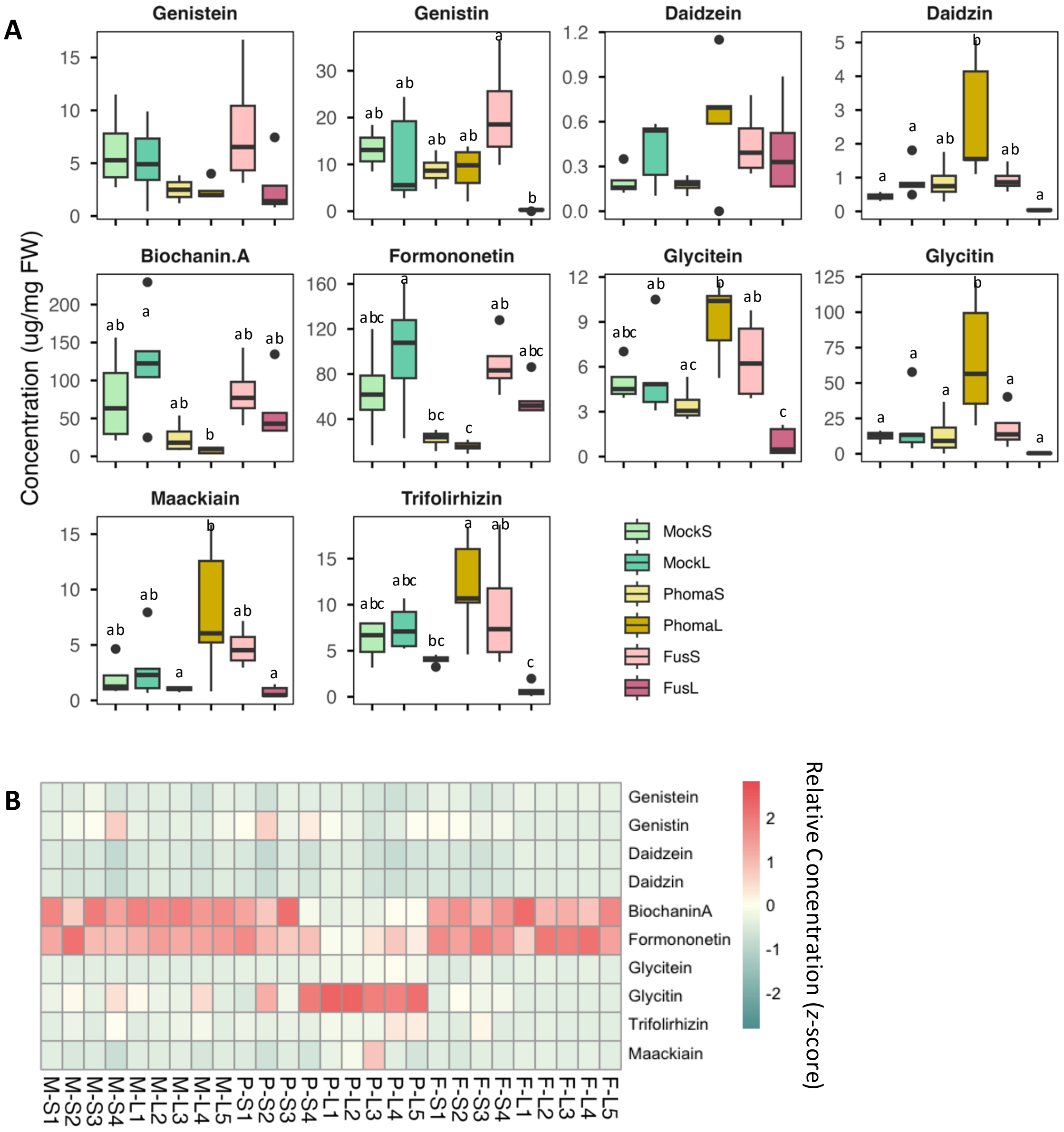


**Supplementary Fig. 1. Quantified isoflavonoids in *T. pratense* root extracts. (A)** Concentrations of compounds quantified against authentic standards. Letters denote significant differences calculated as determined by ANOVA followed by Tukey's HSD test. **(B)** Heatmap of relative concentrations (*z*-scores) normalized within samples.

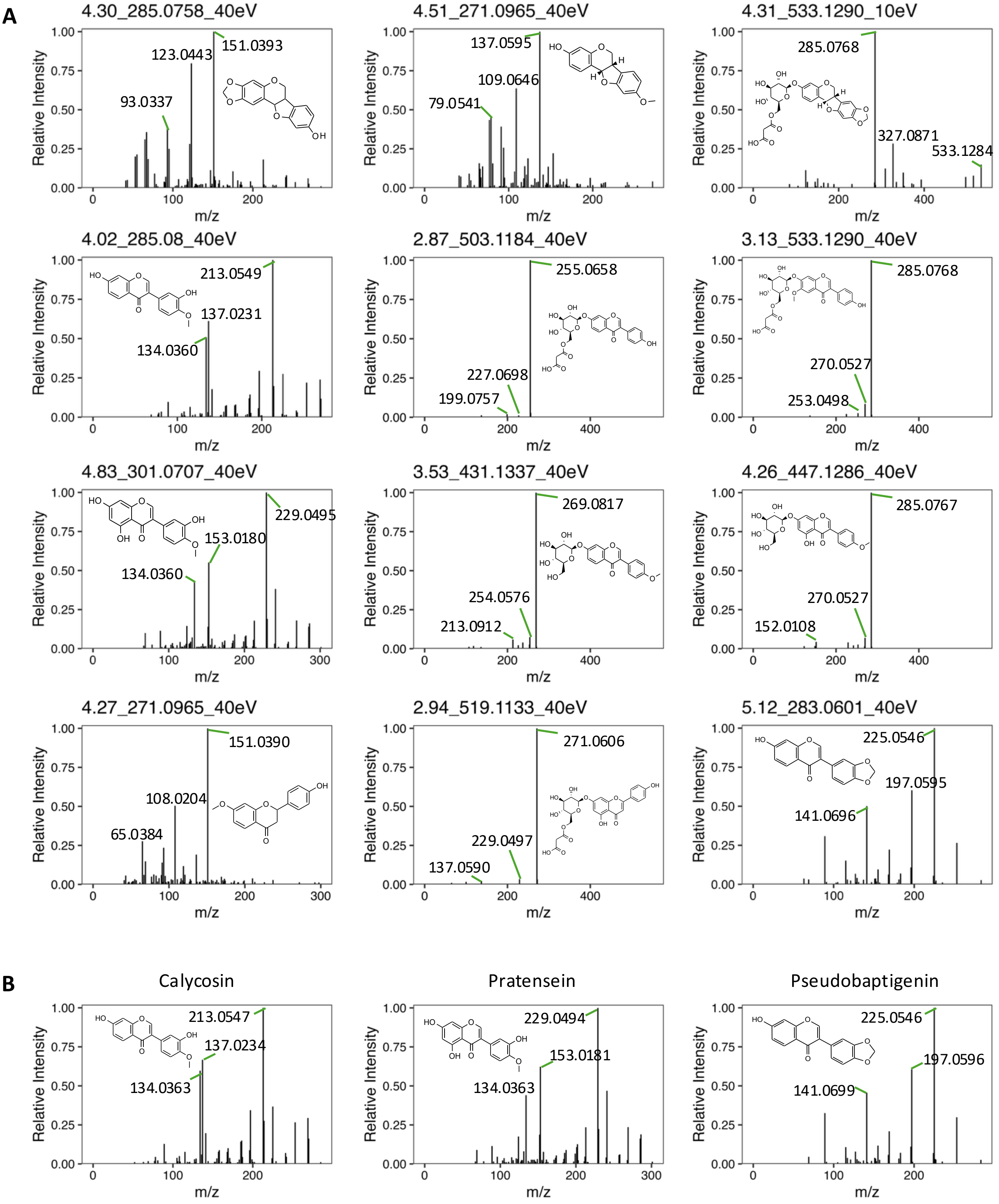


**Supplementary Fig. 2. MS/MS fragmentation spectra of putatively identified compounds in *T. pratense*. (A)** The identities of 12 molecular features were predicted based on their MS/MS spectra. Plots are labelled with *rt*, $[M+H]^+$, collision energy, and putative chemical structure. **(B)** MS/MS spectra of authentic standards (calycosin, pratensein, and pseudobaptigenin) confirm the identities of three putatively identified compounds. All plots are annotated with the *m/z* values of the three most intense peaks.

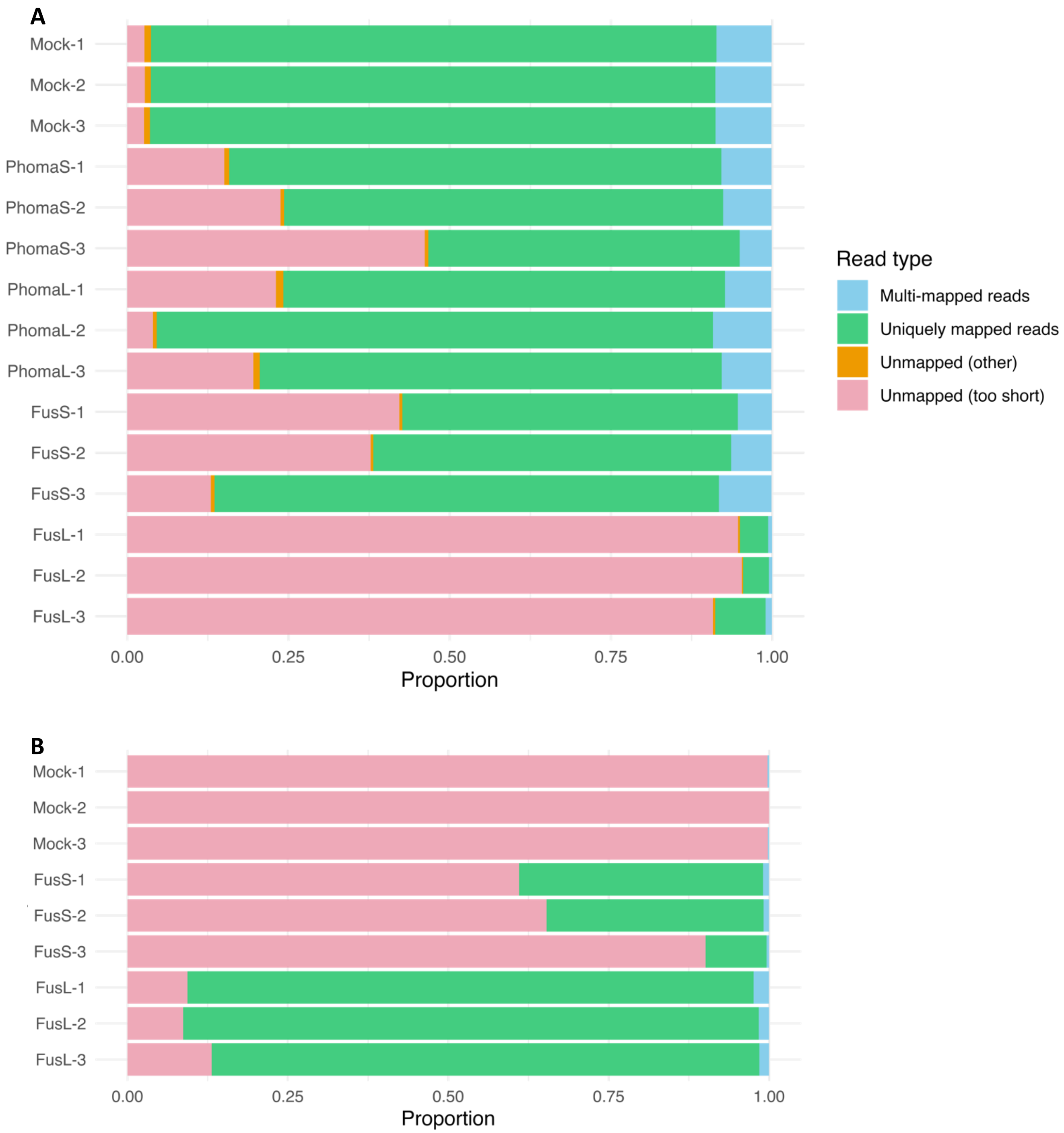


**Supplementary Fig. 3. RNA-Seq STAR alignment statistics.** Visualizing the proportion of multi-mapped, uniquely mapped, and unmapped reads to the **(A)** *T. pratense* NCBI reference genome (RefSeq accession: GCF_020283565.1) and the **(B)** *F. oxysporum* Fo47 reference genome (RefSeq accession: GCF_013085055.1).

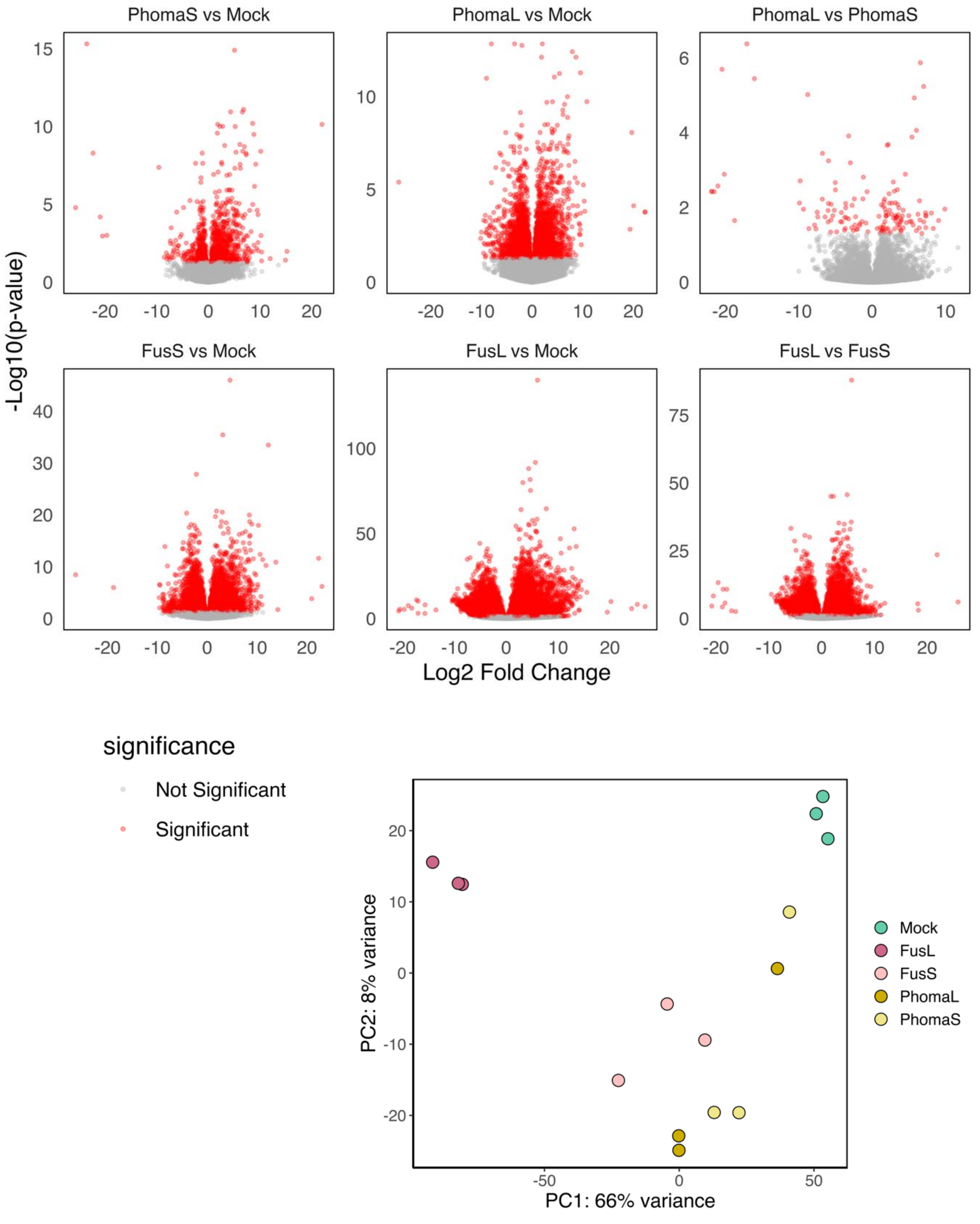


**Supplementary Fig. 4. Transcriptional reprogramming in fungal-infected *T. pratense* roots. (A)** DEG volcano plots of infected tissues relative to mock, or long (7 dpi) relative to short (3 dpi) infection time tissues. Significant genes ($p<0.05$ and absolute log2FC >1) are indicated in red. **(B)** PCA plot of the 15 samples constructed using variance-stabilized gene expression data.

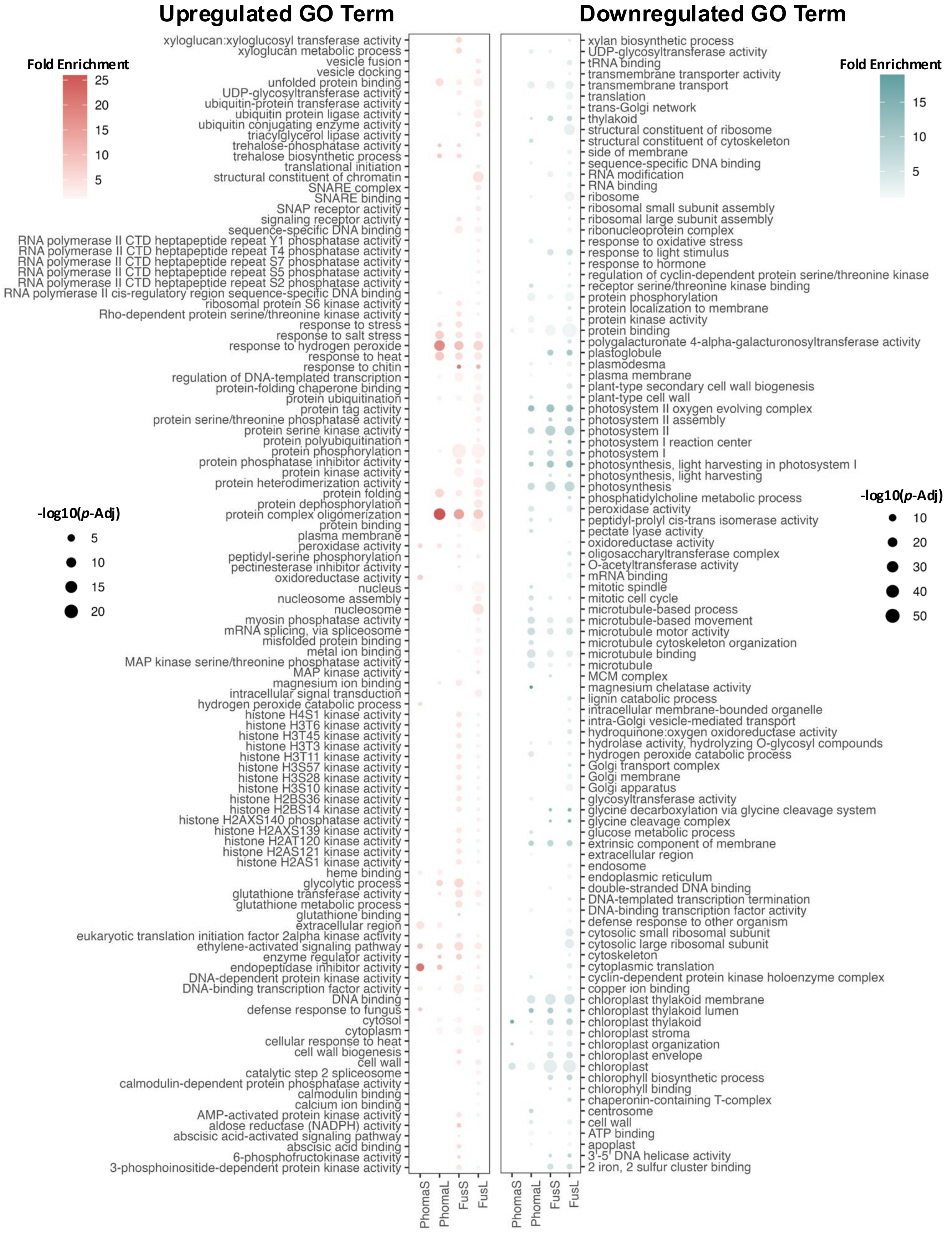


**Supplementary Fig. 5. GO term functional enrichment in response to infection. (A)** Enriched GO terms across up- (red) and downregulated (blue) DEGs (*Benjamini p*-Adj<0.05).

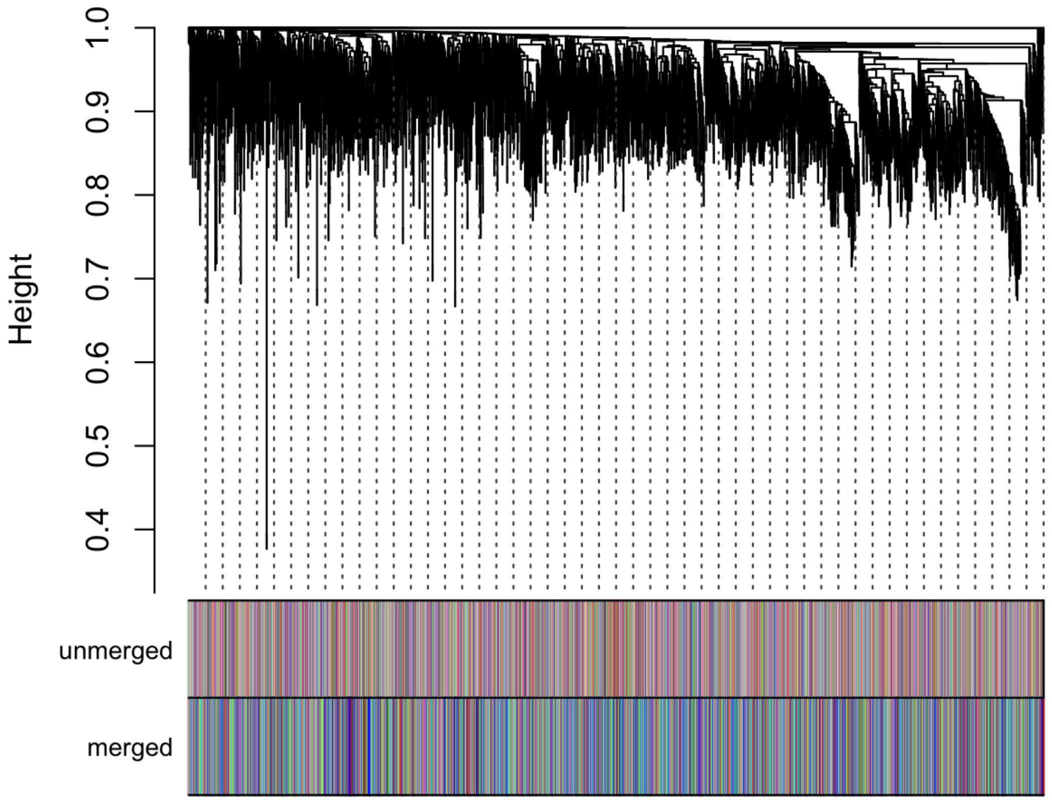


**Supplementary Fig. 6. WGCNA cluster dendrogram.**

**Supplementary Table 1. Classification of 31 molecular features identified in *T. pratense* root extracts.** Compound classes determined based on MS/MS fragmentation matching against natural product fingerprints.

| **Classification** | **Total compounds** |
|---|---|
| Pterocarpan | 3 |
| Pterocarpan malonylglucoside | 1 |
| Hydroxyisoflavone malonylglucoside | 1 |
| *O*-Methylated isoflavone-glucoside | 1 |
| *O*-Methylated hydroxyisoflavone | 3 |
| *O*-Methylated hydroxyisoflavone-glucoside | 2 |
| *O*-Methylated hydroxyisoflavone malonylglucoside | 2 |
| *O*-Methylated dihydroxyisoflavone | 3 |
| *O*-Methylated trihydroxyisoflavone | 4 |
| *O*-Methylated tetrahydroxy isoflavone | 1 |
| Methylenedioxy-hydroxy isoflavone | 1 |
| Methylenedioxy-dihydroxy isoflavone | 2 |
| Trihydroxy-flavone | 2 |
| Dihdroxyflavone malonylglucoside | 1 |
| *O*-Methylated hydroxyflavanone | 2 |
| *O*-Methylated dihydroxyflavanone | 1 |
| *O*-Methylated flavanone-glucoside | 1 |

**Supplementary Table 2. Putatively identified compounds.** Molecular features were assigned putative identities when CSI:FingerID scores exceeded 95%.

| **Top Hit** | **Compound Class** | ***rt* (min)** | **[M+H]+** | **CSI:FingerID scores** |
|---|---|---|---|---|
| Glycitein-derived pterocarpan | Pterocarpan | 4.30 | 285.0758 | 95.06 |
| Sissotrin | *O*-Methylated hydroxyisoflavone-glucoside | 4.26 | 447.1286 | 95.7 |
| 6′-Malonylglycitin | *O*-Methylated hydroxyisoflavone malonylglucoside | 3.13 | 533.129 | 98.1 |
| Ononin | *O*-Methylated isoflavone-glucoside | 3.53 | 431.1337 | 99.7 |
| Pratensein[a] | *O*-Methylated trihydroxyisoflavone | 4.83 | 301.0707 | 95.4 |
| Calycosin[a] | *O*-Methylated dihydroxyisoflavone | 4.02 | 285.0758 | 95.2 |
| 6′-Malonyldaidzin | Hydroxyisoflavone malonylglucoside | 2.87 | 503.1184 | 99.7 |
| Apigenin 7-*O*-malonylglucoside | Dihdroxyflavone malonylglucoside | 2.94 | 519.1133 | 96.1 |
| Methyl-liquiritigenin | *O*-Methylated dihydroxyflavanone | 4.27 | 271.0965 | 97.9 |
| Medicarpin | Pterocarpan | 4.51 | 271.0965 | 98 |
| Trifolirhizin-6''-O-malonate | Pterocarpan malonylglucoside | 4.31 | 533.1290 | 99.2 |
| Pseudobaptigenin[a] | Methylenedioxy-hydroxy isoflavone | 5.12 | 283.0601 | 90.1 |

[a] MS2 fragmentation spectra further confirmed by comparison to authentic standards

**Supplementary Table 3. Most highly expressed P450s across all root samples.** Mean and variance were calculated using variance-stabilized gene expression data across all samples.

| geneID | Enzyme | Mean expression | Variance |
|---|---|---|---|
| LOC123918472 | IFS | 14.73 | 1.92 |
| LOC123884908 | I3'H | 13.69 | 0.01 |
| LOC123894475 | P1[a] | 13.66 | 1.93 |
| LOC123884134 | I3'H | 13.42 | 0.11 |
| LOC123906372 | | 12.85 | 2.43 |
| LOC123892439 | | 12.52 | 4.36 |
| LOC123894994 | P5[a] | 12.38 | 0.96 |
| LOC123906491 | | 12.24 | 2.03 |
| LOC123900153 | P4[a] | 12.15 | 0.82 |
| LOC123919903 | | 12.13 | 0.29 |

[a] Candidate genes screened in this study.

**Supplementary Table 4.** Percent identity matrix of five selected candidate P450s generated by ClustalW.

| | P5 | P2 | P4 | P1 | P3 |
|---|---|---|---|---|---|
| **P5 (LOC123894994)** | 100.00 | 20.53 | 19.70 | 18.32 | 19.82 |
| **P2 (LOC123909586)** | 20.53 | 100.00 | 23.55 | 26.04 | 26.62 |
| **P4 (LOC123900153)** | 19.70 | 23.55 | 100.00 | 26.95 | 28.99 |
| **P1 (LOC123894475)** | 18.32 | 26.04 | 26.95 | 100.00 | 32.24 |
| **P3 (LOC123906377)** | 19.82 | 26.62 | 28.99 | 32.24 | 100.00 |

**Supplementary Table 5.** Percent identity matrix of the TpPbS protein sequence compared to other P450s of interest.

| | CYP719A24 | CYP719A5 | CYP719A14 | CYP719A13 | CYP719A2 | CYP81Q1 | CYP81Q2 | CYP76F112 | CYP76F1 | CYP76F319 | CYP76F71 | CYP76F70 |
|---|---|---|---|---|---|---|---|---|---|---|---|---|
| **CYP719A24** | 100 | 55.04 | 52.08 | 55.44 | 55.35 | 26.54 | 26.54 | 26.16 | 23.06 | 23.75 | 23.54 | 22.92 |
| **CYP719A5** | 55.04 | 100 | 79.14 | 61.32 | 62.66 | 25.16 | 25.38 | 27.29 | 26.71 | 26.33 | 26.06 | 26.48 |
| **CYP719A14** | 52.08 | 79.14 | 100 | 59.39 | 61.44 | 22.86 | 22.86 | 28.3 | 26.69 | 26.11 | 25.89 | 26.32 |
| **CYP719A13** | 55.44 | 61.32 | 59.39 | 100 | 78.34 | 25.48 | 25.05 | 26.68 | 25.83 | 25.1 | 25.31 | 25.31 |
| **CYP719A2** | 55.35 | 62.66 | 61.44 | 78.34 | 100 | 25.32 | 24.89 | 27.23 | 26.64 | 27.27 | 26.43 | 26.64 |
| **CYP81Q1** | 26.54 | 25.16 | 22.86 | 25.48 | 25.32 | 100 | 95.85 | 30.04 | 29.46 | 29.81 | 30.45 | 30.04 |
| **CYP81Q2** | 26.54 | 25.38 | 22.86 | 25.05 | 24.89 | 95.85 | 100 | 29.42 | 29.67 | 30.02 | 30.25 | 29.84 |
| **CYP76F112** | 26.16 | 27.29 | 28.3 | 26.68 | 27.23 | 30.04 | 29.42 | 100 | 51.41 | 53.32 | 51.2 | 52.4 |
| **CYP76F1** | 23.06 | 26.71 | 26.69 | 25.83 | 26.64 | 29.46 | 29.67 | 51.41 | 100 | 84.89 | 82.18 | 83.96 |
| **CYP76F319** | 23.75 | 26.33 | 26.11 | 25.1 | 27.27 | 29.81 | 30.02 | 53.32 | 84.89 | 100 | 87.15 | 89.72 |
| **CYP76F71** | 23.54 | 26.06 | 25.89 | 25.31 | 26.43 | 30.45 | 30.25 | 51.2 | 82.18 | 87.15 | 100 | 93.32 |
| **CYP76F70** | 22.92 | 26.48 | 26.32 | 25.31 | 26.64 | 30.04 | 29.84 | 52.4 | 83.96 | 89.72 | 93.32 | 100 |
